# Protective efficacy of an orf virus-vector encoding the hemmagglutinin and the nucleoprotein of influenza A virus in swine

**DOI:** 10.1101/2021.04.19.440556

**Authors:** Lok R. Joshi, David Knudsen, Pablo Pineyro, Santhosh Dhakal, Gourapura J. Renukaradhya, Diego G. Diel

## Abstract

Swine influenza is a highly contagious respiratory disease of pigs caused by influenza A viruses (IAV-S). IAV-S causes significant economic losses to the swine industry and poses constant challenges to public health due to its zoonotic potential. Thus effective IAV-S vaccines are highly desirable and would benefit both animal and human health. Here, we developed two recombinant orf viruses, expressing the hemagglutinin (HA) gene (OV-HA) or both the HA and the nucleoprotein (NP) genes of IAV-S (OV-HA-NP). The immunogenicity and protective efficacy of these two recombinant viruses were evaluated in pigs. Both OV-HA and OV-HA-NP recombinants elicited robust virus neutralizing antibody response in pigs. Notably, although both recombinant viruses elicited IAV-S-specific T-cell responses, the frequency of IAV-S specific proliferating T cells secreting IFN-γ upon re-stimulation was higher in OV-HA-NP-immunized animals than in the OV-HA group. Importantly, IgG1/IgG2 isotype ELISAs revealed that immunization with OV-HA induced Th2-biased immune responses, whereas immunization with OV-HA-NP virus resulted in a Th1-biased immune response. While pigs immunized with either OV-HA or OV-HA-NP were protected when compared to non-immunized controls, immunization with OV-HA-NP resulted in better protective efficacy as evidenced by reduced virus shedding in nasal secretions and reduced viral load in the lung. This study demonstrates the potential of ORFV-based vector for control of swine influenza virus in swine.

**Importance:** Effective influenza A virus (IAV-S) vaccines capable of providing robust protection to genetically diverse IAV-S in swine are lacking. Here, we explored the potential of orf virus based vectors expressing the hemagglutining (HA) or both the HA and the nucleoprotein (NP) genes of influena A virus (IAV-S) in eliciting protection against IAV-S in pigs. We observed that both recombinant viruses elicited IAV-S-specific humoral and cell-mediated immune responses in pigs. Addition of the NP and co-expression of this protein with HA, another major influenza protective antigen, resulted in higher T cell responses which presumably led to better protection in OV-HA-NP immunized animals, as evidenced by lower levels of virus shedding and viral load in lungs. This study highlights the the potential of ORFV as a vector platform for vaccine delivery against IAV-S. Results here provide the foundation for future development of broadly protective ORFV-based vectors for IAV-S for use in swine.

## Introduction

Swine influenza is a highly contagious respiratory disease of pigs caused by influenza A viruses in swine (IAV-S). IAV-S is an enveloped, single stranded RNA virus of the family *Orthomyxoviridae*. The IAV-S genome consists of eight single-stranded negative-sense RNA segments encoding four structural (HA, NA, NP and M) and four non-structural (PB1, PB2, PA and NS) proteins. Influenza viruses are classified into subtypes based on the antigenicity of hemagglutinin (HA) and neuraminidase (NA) proteins present on the surface of the virus. There are three recognized subtypes of IAV-S that are currently circulating in the US: H1N1, H1N2 and H3N2 (1). The H1N1 subtype is the major subtype that has been prevalent in the US swine population for several decades; however, recent epidemiological data suggests an increasing incidence of H1N2 and H3N2 IAV-S subtypes (2, 3). IAV-S causes acute respiratory disease in pigs resulting in high morbidity (up to 100%). The mortality rate is usually low (1-4%) with most infected animals recovering within 3-7 days of infection (4, 5). The median yearly herd prevalence of IAV-S reported in the US is approximately 28%, but it can reach up to 57% in winter and spring months (6). IAV-S results in significant economic losses to the swine industry mainly due to weight loss, increased time to market, costs associated with treatment of secondary bacterial infections and mortality. This makes IAV-S one of the top three health challenges to the swine industry affecting pigs in all phases of production (7, 8). In addition to IAV-S, pigs are also susceptible to infection with avian and human IAVs thereby providing a niche for genetic reassortment between avian/human or swine influenza viruses. This poses a major threat for emergence of new subtypes as well as increases the risk of zoonotic transmission of IAVs. Therefore, effective prevention and control measures for IAV infections in swine have direct impacts on both animal and human health.

Currently, most available IAV-S vaccines are based on whole inactivated virus (WIV). However, these vaccines have not been able to effectively control IAV in swine and in some cases vaccine associated enhanced respiratory disease has been observed when there is an antigenic mismatch between vaccine strain and infecting strain (9). A live-attenuated influenza virus (LAIV) vaccine based on a virus containing a deletion of the NS1 gene, has been recently licensed for use in pigs in the US and may overcome some of the drawbacks of WIV vaccines (10). However, LAIV vaccines have the potential to reassort with the endemic viruses potentially resulting in new influenza virus variants. Indeed, novel variants that arose from reassortment between the vaccine virus and endemic field strains have been recently reported (11). These observations highlight the need for safer and more efficaceous IAV-S vaccine candidates. Here we investigated the potential of vectored vaccine candidates based on the parapox orf virus (ORFV) in controlling IAV-S infection in pigs.

Orf virus (ORFV) belongs to genus *Parapoxvirus* within the family *Poxviridae* (12) and is a ubiquitous virus that primarily causes a self-limiting mucocutaneous infection in sheep, goats and wild ruminants (13, 14). ORFV contains a double-stranded DNA genome with approximately 138 kbp in length and encodes 131 putative genes, including several with immunomodulatory (IMP) functions (15). Given ORFV IMP properties, the virus has long been used as a preventive and therapeutic agent in veterinary medicine (16, 17). Additionally, the potential of ORFV as a vaccine delivery platform against several viral diseases in permissive and non-permissive animal species has been explored by us and others (18–24). ORFV based vectored-vaccine candidates have been shown to induce protective immunity against pseudorabies virus (PRV), classical swine fever virus (CSFV) and porcine epidemic diarrhea virus (PEDV) (22, 25–27). Among the features that make ORFV a promising viral vector for vaccine delivery in swine are : (i) its restricted host range, (ii) its ability to induce both humoral and cellular immune response (22, 28), (iii) its tropism which is restricted to skin keratinocytes with no evidence of systemic dissemination, (iv) lack of vector-specific neutralizing antibodies which allows efficient prime-boost strategies using the same vector constructs (29, 30), and (v) its large genome size with the presence of several non-essential genes, which can be manipulated without severely impacting virus replication. Additionally, ORFV encodes several genes with well-characterized immunomodulatory properties. These include a homologue of interleukin 10 (IL-10) (31), a chemokine binding protein (CBP) (32), an inhibitor of granulocyte-monocyte colony stimulating factor GMC-CSF) (33), an interferon resistance gene (VIR) (34), a homologue of vascular endothelial growth factor (VGEF) (35), and inhibitors of nuclear-factor kappa-B (NF-ƙB) signaling pathway (36–39). The presence of these well-characterized immunomodulatory proteins allowed us to rationally engineer ORFV-based vectors with enhanced safety and immunogenicity profile for use in livestock species, including swine (22–24).

Here we assessed the immunogenicity and protective efficacy of recombinant ORFV vectors expressing the HA protein alone or the HA and the nucleoprotein (NP) of IAV-S. While the HA protein contains immunodominant epitopes recognized by neutralizing antibodies (40, 41), the NP protein contains highly conserved immunodominant T-cell epitopes (42). We investigated whether co-expression of HA and NP would enhance the protective efficacy against IAV-S following intranasal challenge infection in pigs.

## Results

### Construction of ORFV recombinants

The OV-HA recombinant virus was obtained by inserting the full-length HA gene of IAV-S (H1N1) into ORFV121 locus by homologous recombination between a transfer plasmid pUC57-121LR-SIV-HA-loxp-GFP and the parental ORFV strain IA82 (Fig 1A). The OV-HA-NP recombinant virus was obtained by inserting the full-length HA gene into ORFV121 locus and the NP gene into ORFV127 locus. The wild type ORFV strain IA82 was used to generate the OV-HA virus which served as a parental virus for generation of the OV-HA-NP recombinant (Fig 1B). Expression of HA was driven by the vaccina virus (VACV) I1L promoter (43) and expression of the NP gene was driven by the VACV vv7.5 promoter (44). After infection with the parental virus and transfection with the recombination plasmid, the recombinant viruses were obtained and selected. Several rounds of plaque assays were performed to obtain purified recombinant viruses. Once the recombinant viruses were purified and verified by PCR, the marker gene encoding for the green fluorescent protein (GFP) was removed by using the Cre recombinase system. Whole-genome sequencing of plaque purified recombinant viruses was performed after Cre recombinase treatment. Sequencing results confirmed the integrity and identity of ORFV sequences, demonstrated the presence of HA gene, deletion of ORFV121 in OV-HA construct, the presence of HA and NP genes and deletion of ORFV121 and ORFV127 genes in OV-HA-NP construct.

**Figure 1.**
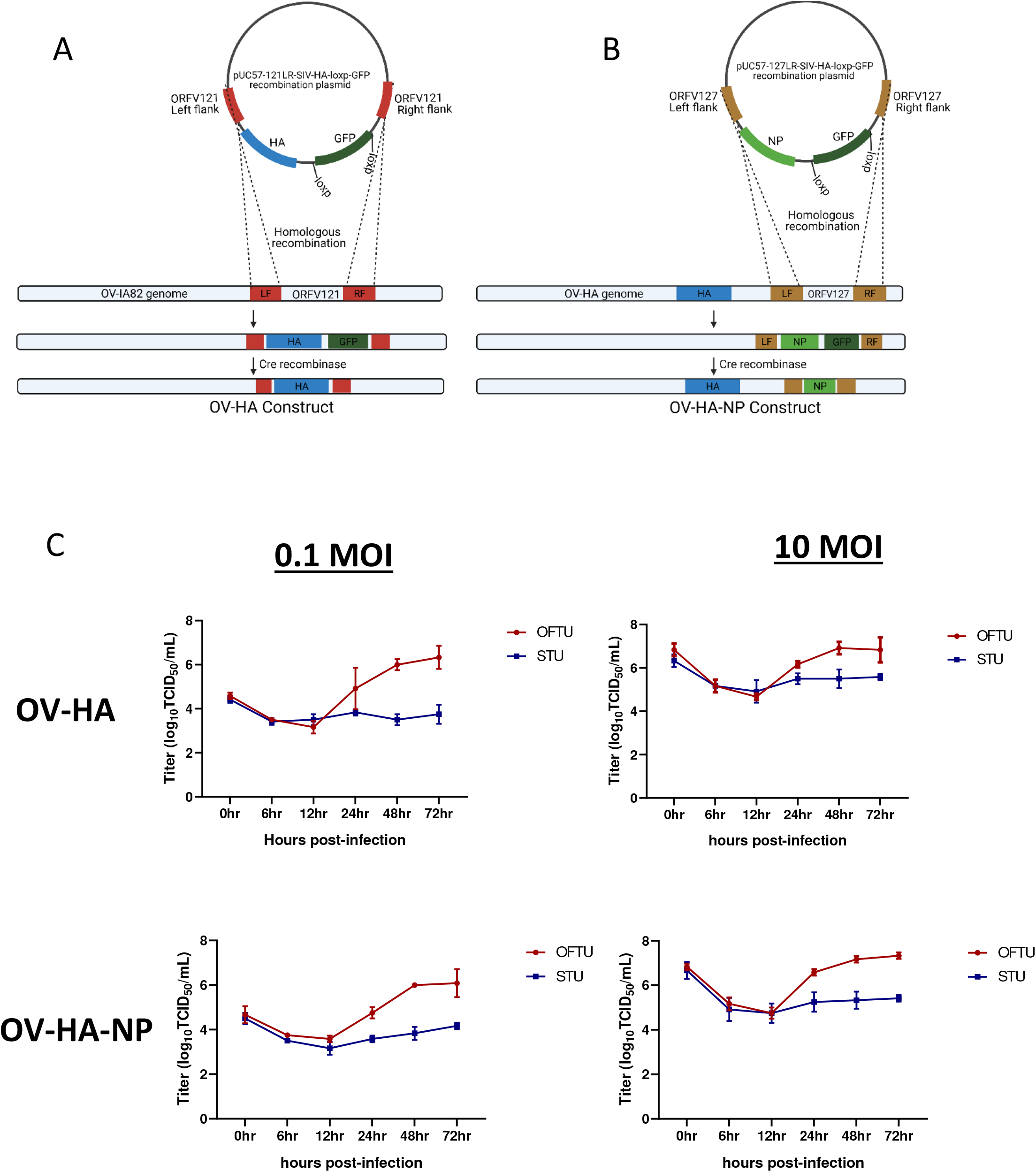
Construction of ORFV recombinants and their replication kinetics. **(A)** Schematic representation of homologous recombination between pUC57-121LR-SIV-HA-loxp-GFP plasmid and ORFV-IA82 genome. The recombinant virus was treated with Cre recombinase to remove GFP marker gene to obtain markerless OV-HA construct. **(B)** Schematic representation of homologous recombination between pUC57-127LR-SIV-NP-loxp-GFP plasmid and OV-HA genome. The recombinant virus was treated with Cre recombinase to obtain markerless OV-HA-NP construct. **(C)** Multi-step (0.1 MOI) and single step (10 MOI) growth curve of OV-HA and OV-HA-NP. OFTu or STU cells were infected with OV-HA and-HA-NP recombinants and virus titers were calculated at 0, 6, 12, 24, 48 and 72 hours post-infection. Error bars represent SEM calculated based on three independent experiments.

### Replication kinetics of OV-HA and OV-HA-NP viruses *in vitro*

Replication properties of both recombinant viruses (OV-HA and OV-HA-NP) were assessed *in vitro* in primary ovine fetal turbinate cells (OFTu) and primary swine turbinate cells (STU) using one-step and multi-step growth curves (Fig 1C). Cells were infected with an MOI of 0.1 or 10 and cell lysates were harvested at 6, 12, 24, 48, 72 hours post-infection. Both recombinants replicated efficiently in natural host OFTu cells. However, replication of OV-HA and OV-HA-NP viruses was markdely impaired in the STU cells (Fig 1C), which increases the safety profile of the vector for use in pigs.

### Expression of heterologous proteins by OV-HA and OV-HA-NP recombinant viruses

Expression of the HA protein and NP proteins by OV-HA and/or OV-HA-NP viruses was confirmed by immunofluorescence assay (IFA) and flow-cytometry. As shown in the figure 2A, OV-HA recombinant expressed high levels of HA and OV-HA-NP recombinant expressed high levels of HA and NP proteins (Fig 2A). Expression of HA and NP were also confirmed by flow cytometry (Fig 2C). The IFA was also performed in non-permeabilized cells. Both HA and NP proteins were detected in non-permeabilied cells; however, the levels of protein detected were slightly lower than in permeabilized cells (Fig 2B). As expected this decrease was more evident for NP protein than for the HA protein. These findings suggest that while a great proportion of the HA protein expressed by both OV-HA and OV-HA-NP recombinant viruses localizes to the cell surface, and expression of the NP protein is mostly confined to the intracellular compartment.

**Figure 2.**
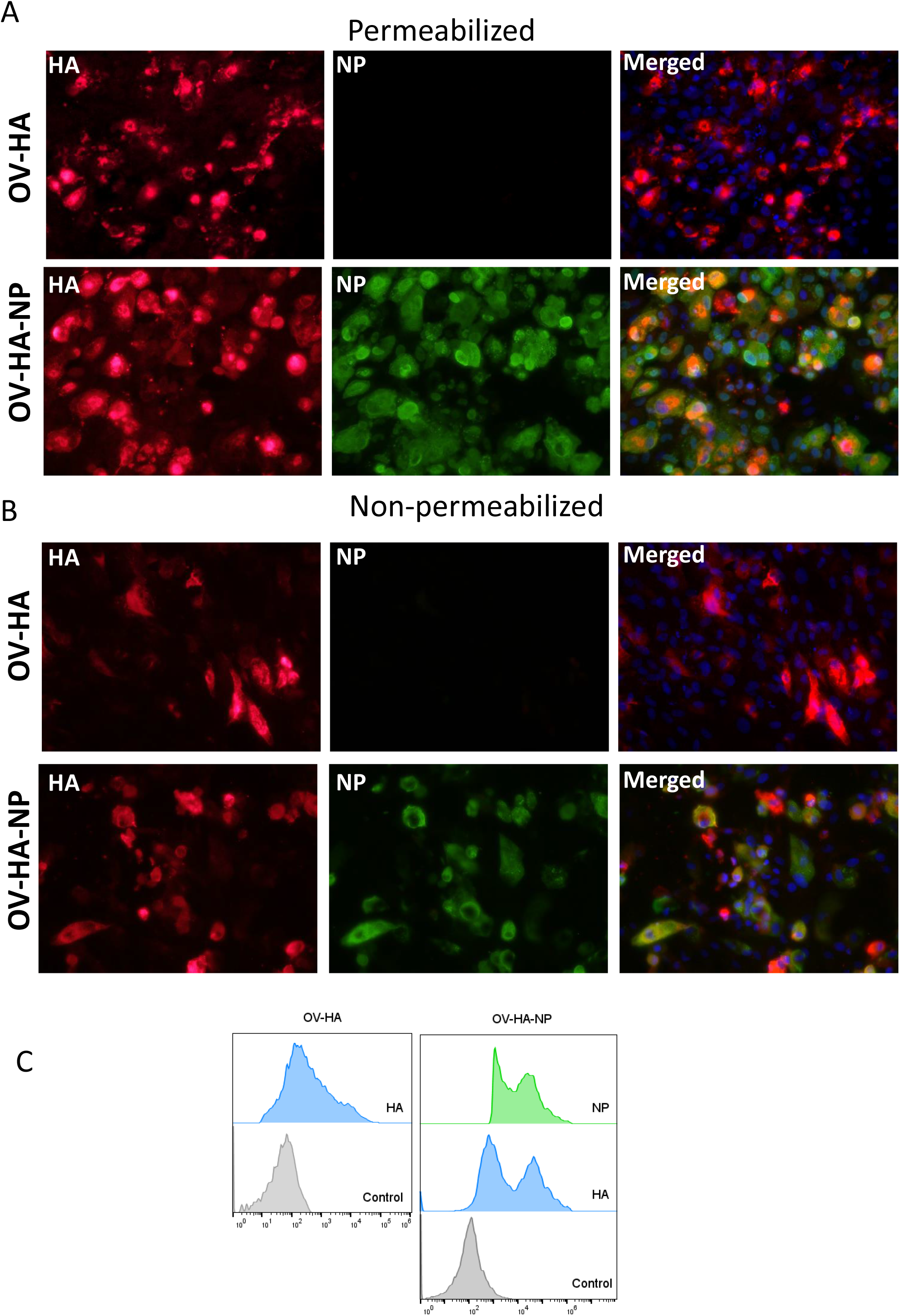
Expression of heterologous proteins by ORFV recombinants. **(A)** Immunofluorescence assay in permeabilized OFTu cells. Upper panel shows expression of HA protein and absence of NP protein in OV-HA recombinant. Lower panel shows expression of HA and NP protein by OV-HA-NP recombinant. **(B)** Immunofluorescence assay performed in non-permeabilized OFTu cells. Upper panel shows expression of HA by OV-HA recombinant and lower panel shows expression of HA and NP by OV-HA-NP recombinant. Blue fluorescence in merged images in panel A and B indicates nuclear staining by DAPI. **(C)** Expression of heterologous proteins by ORFV recombinants assessed by flow-cytometry. OFTu cells were infected with OV-HA, OV-HA-NP or Wild-type OV-IA82 as negative control. Infected cells were collected 48 hours post-infection, fixed and then stained with appropriate antibodies for flow cytometric analysis.

### Immunigenicity of OV-HA and OV-HA-NP in pigs

To assess the immunogenicity of OV-HA and OV-HA-NP, 4-week old, IAV-S negative, weaned piglets were immunized intramuscularly with two doses of OV-HA and OV-HA-NP at a 21 day interval (Fig 3A; Table 1). Antibody response were evaluated using virus neutralization (VN) and hemagglutination inhibition (HI) assays. One week after the first immunization, neutralizing antibodies were detected in both vaccinated groups, however the levels were significantly higher in OV-HA-NP vaccinated animals (Fig 3B). An anamnestic increase in neutralizing antibody titers was seen in both vaccinated groups one week after the boost immunization (28 days post-immunization). After the booster immunization all animals maintained high level of neutralizing antibody levels untill the end of the experiment (42 dpi, Fig. 3B).

**Figure 3.**
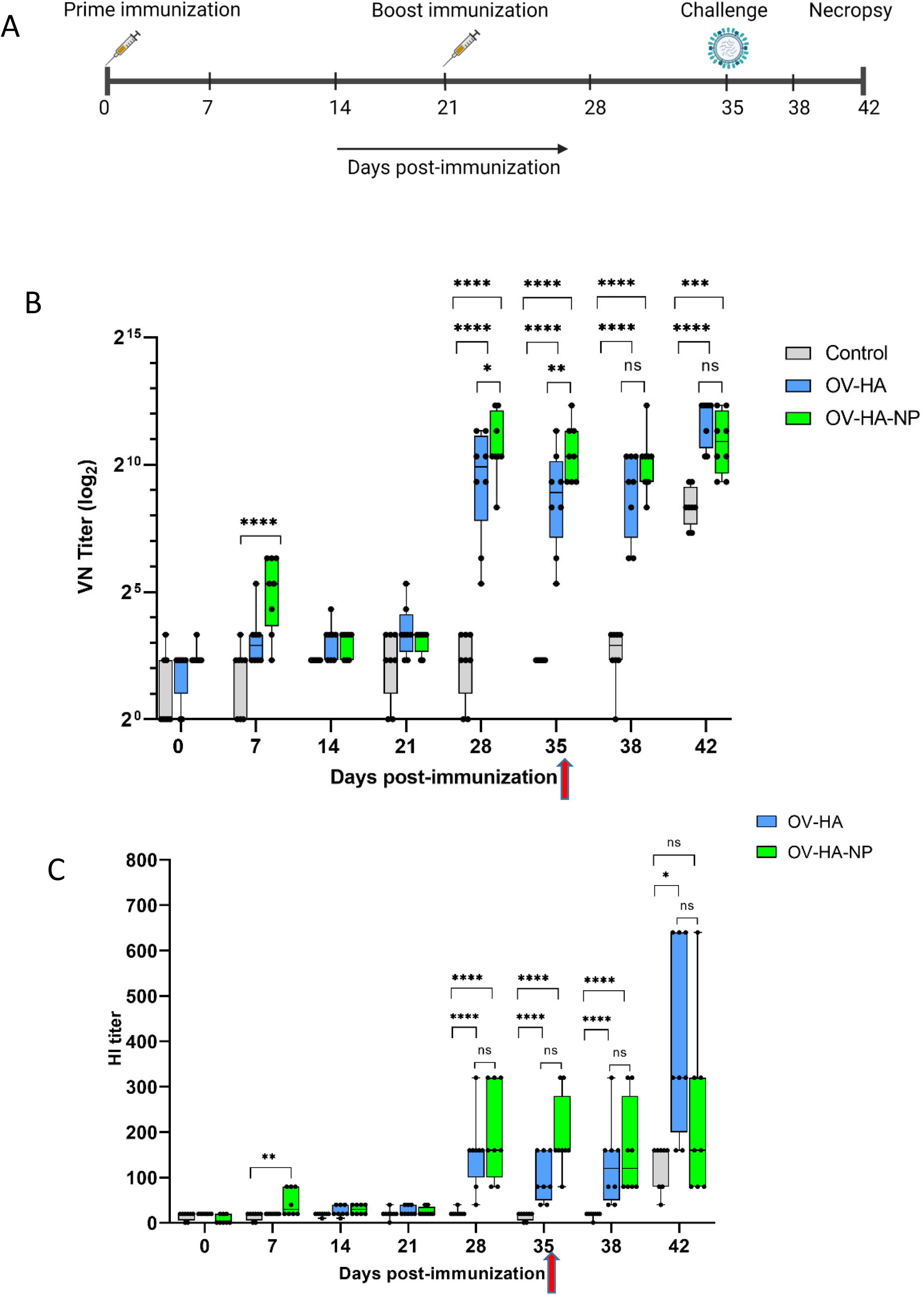
Immunization-challenge experiment design and humoral response to immunization. **(A)** A timeline of immunization-challenge experiment. **(B)** IAV-S specific neutralizing antibody response elicited by immunization with OV-HA and OV-HA-NP. **(C)** IAV-S specific humoral immune response induced by OV-HA and OV-HA-NP assessed by hemagglutination inhibition (HI) assay. Red arrow heads represent the day of challenge. The error bars represent SEM. VN titer shown in logarithmic scale for effective visualization. HI titer shown in liner scale. P-values: *\*P < 0*.*05, **P < 0*.*01, ***P < 0*.*001, ****P < 0*.*0001*.

**Table 1.** Experimental design for immunization-challenge study

| Group | Immunization | Number of animals | Immunization Days | Immunization Route | Challenge | Challenge Dose (Route) |
| --- | --- | --- | --- | --- | --- | --- |
| 1 | Control | 8 | 0, 21 | IM | H1N1<br>A/Swine/OH/24366/2007 | $6 \times 10^6$ TCID<sub>50</sub><br>(Intranasal) |
| 2 | OV-HA | 8 | 0, 21 | IM | H1N1<br>A/Swine/OH/24366/2007 | $6 \times 10^6$ TCID<sub>50</sub><br>(Intranasal) |
| 3 | OV-HA-NP | 8 | 0, 21 | IM | H1N1<br>A/Swine/OH/24366/2007 | $6 \times 10^6$ TCID<sub>50</sub><br>(Intranasal) |

Serological responses were also measured using an hemagglutination inhibition (HI) assay. The presence of HI antibodies were detected in OV-HA-NP group on day 7 pi. Similar to the VN results, an anamnestic increase in HI antibody titers was observed one week after the booster immunization in both groups (Fig 3C). Interestingly, the HI titers in the OV-HA group increased significantly after challenge, which is more evident a week after challenge (42 dpi). Such anamnestic increase in HI titers was not seen in OV-HA-NP-immunized animals, suggesting enhanced protection from IAV-S challenge in this group (Fig 3C). Overall, these results demonstrate that immunization with OV-HA and OV-HA-NP viruses elicited high IAV-S specific neutralizing and HI antibody responses in immunized pigs.

### IAV-S-specific IgG isotype responses elicited by immunization with OV-HA and OV-HA-NP viruses

IAV-S specific IgG responses were measured using a whole virus ELISA. Low levels of IAV-S-specific total IgG antibodies were detected in OV-HA and OV-HA-NP immunized groups on 21 days pi (Fig 4A). Similar to VN and HI assay, significantly higher levels of IgG antibodies was observed a week following the boost immunization (day 28 pi). Thereafter consistently higher levels of IgG were detected in serum of both OV-HA and OV-HA-NP immunized groups until the end of the experiment (Fig 4A). As expected, expression and delivery of the NP by the OV-HA-NP recombinant virus elicited higher levels of IgG antibodies in immunized pigs after the booster immunization on day 21 pi when compared to those observed in OV-HA-immunized animals (*P < 0*.*0001*, Fig 4A).

**Figure 4.**
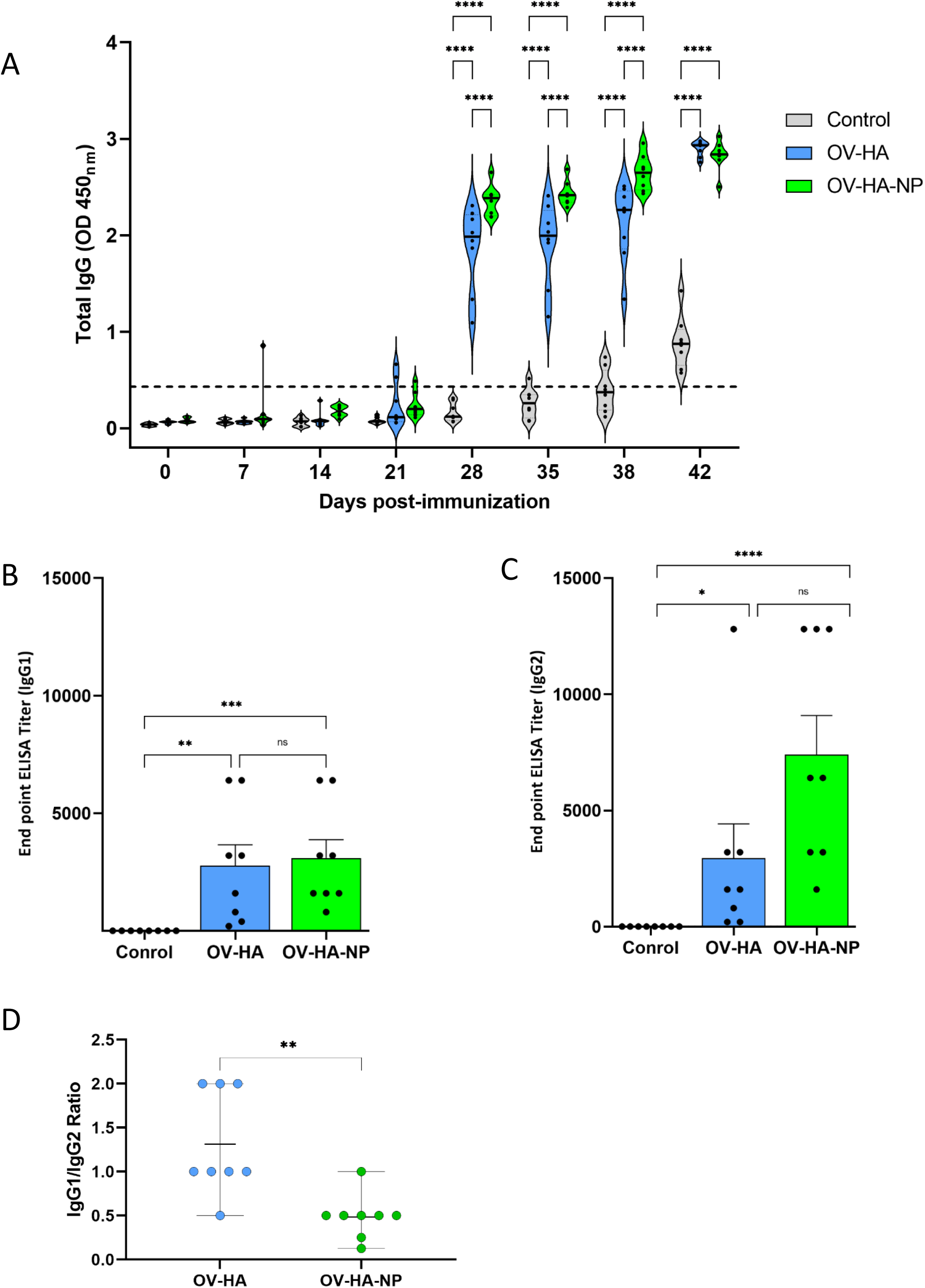
IAV-S specific IgG responses to immunization. **(A)** Total serum IgG level elicited by OV-HA and OV-HA-NP immunization at various time points were assessed by ELISA. Isotype ELISA demonstrating endpoint titers elicited by immunization at 35 days pi in serum was assessed for detecting specific **(B)** IgG1 and **(C)** IgG2 antibodies. **(D)** IgG1/IgG2 ratio in immunized animals was calculated. Each dot represents IgG1/IgG2 ratio of an individual animal. Middle bar represents mean ratio and upper and lower bars represent range. P-values: *\*P < 0*.*05, **P < 0*.*01, ***P < 0*.*001, ****P < 0*.*0001*.

The endpoint titer of IgG1 and IgG2 isotype antibodies elicited by immunization with OV-HA and OV-HA-NP were determined by an isotype ELISA performed on serum samples collected on 35 days pi. Immunization with OV-HA and OV-HA-NP viruses elicited similar levels of IgG1 response, however, significantly higher titers of IgG2 antibodies were detected in OV-HA-NP-immunized animals when compared to IgG2 titers detected in OV-HA-immunized animals (Fig 4B and 4C). The ratio of Th2-associated IgG1 isotype and Th1-associated IgG2 isotype (IgG1/IgG2 ratio) calculated based on the endpoint titers detected in each group was 1.31 (i.e. > 1) for the OV-HA group and 0. 48 (i.e. <1) for the OV-HA-NP group (Fig 4D). The IgG1/IgG2 ratio in OV-HA-NP group was significantly lower than in the OV-HA group (P = 0.0048, Mann-Whitney test). Together these results suggest that the immune response in OV-HA group is mostly Th2 biased. In contrast, the immune response was Th1 biased on the OV-HA-NP group as indicated by higher levels of IgG2 antibodies in the serum of OV-HA-NP immunized animals.

### Cellular immune responses elicited by immunization with OV-HA and OV-HA-NP

IAV-S-specific T-cell responses elicited by immunization with OV-HA and OV-HA-NP viruses was assessed on peripheral blood mononuclear cells (PBMCs) collected on 35 days pi (pre-challenge infection). The frequency of different T-cell subsets secreting IFN-γ following re-stimulation with IAV-S was measured using intracellular cytokine staining (ICS) assays. Upon singlet selection, live/dead cell discrimination, IFN-γ expression by different T-cell subsets including total T-cells (CD3+), CD4+ T-cells (CD3+/CD4+), CD8+ T-cells (CD3+/CD4-CD8+), double positives (CD3+/CD4+/CD8+) and double negative T-cells (CD3+/CD4-/CD8-) were assessed. Animals immunized with either OV-HA or OV-HA-NP had significantly higher percentage of CD3^+^ T-cells secreting IFN-γ when compared to the non-immunized control animals (Fig 5A). Notably, within the vaccinated animals, OV-HA-NP group presented a significantly higher frequency of IFN-γ secreting CD3^+^ T-cells than the OV-HA group (*P=0*.*0055*). The animals in the OV-HA-NP group presented higher frequency of IFN-γ secreting CD3+/CD4+ T-cells, however, the differences between the groups was not statistically significant. Both immunized groups presented increased frequencies of CD3+/CD8+, CD3+/CD4+/CD8+ (double positives) and CD3+/CD4-/CD8-(double negative) IFN-γ secreting T-cell subsets when compared to the control sham-immunized group (Fig 5A).

**Figure 5.**
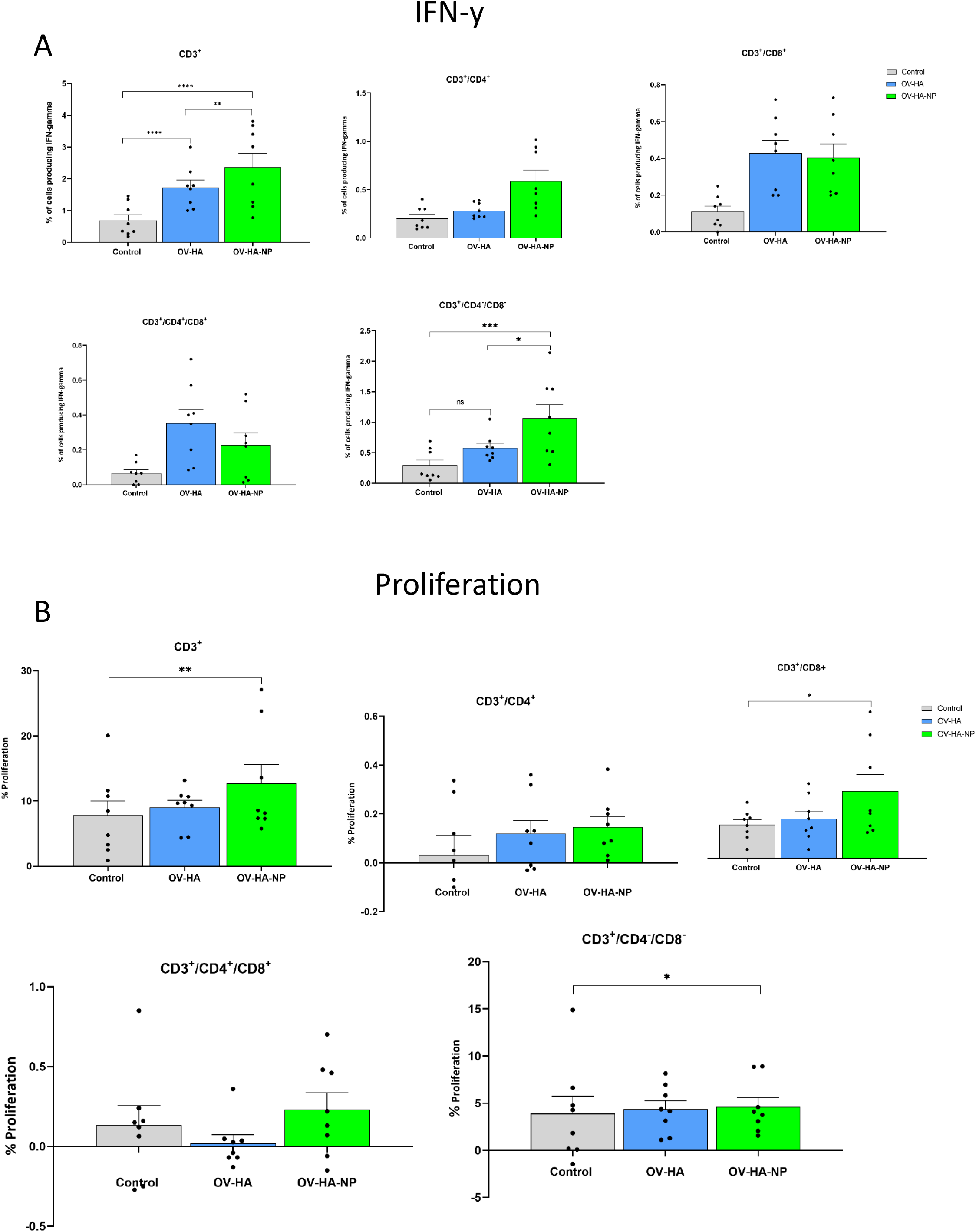
T-cell immune response to immunization. PBMCs isolated from pigs at 35 dpi following recall stimulation with inactivated IAV-S were analysed for: **(A)** IFN-γ production by different T-cell subsets measured by flow cytometry assay; and **(B)** T-cells proliferation by CFSE dilution assay. Data represents group means and error bars represent SEM. P-values: *\*P < 0*.*05, **P < 0*.*01, ***P < 0*.*001, ****P < 0*.*0001*.

IAV-S-specific T-cell responses were also evaluated by the carboxyfluorescein succinimidyl ester (CFSE) dilution assay to determine the specific T-cell subsets proliferating upon re-stimulation of PBMCs with inactivated IAV-S. As described above for the IFN-γ ICS, upon singlet selection and dead cell exclusion, proliferation by the major swine T-cell subsets was evaluated (Fig 5B). While proliferation of CD3+ T-cell subset was observed in animals immunized with OV-HA or OV-HA-NP, significant proliferation of CD3+ T-cells was observed in OV-HA-NP group upon recall stimulation (*P*=0.0095; Fig 5B). Additionally, a significant increase in the proliferation of CD3+/CD8+ T-cell subset was observed in the OV-HA-NP group (*P=0*.*0217*, Fig 5B). An increase in proliferation of CD3+/CD4+ T-cells was also observed (Fig 5B); however, the differences between the treatment groups were not statistically significant (Fig 5B). Overall, these results show that both OV-HA and OV-HA-NP group were able to induce IAV-S-specific T-cell responses in the immunized animals. As expected, T-cell responses elicited by immunization with the OV-HA-NP construct was higher than those observed in animals immunized with the single gene OV-HA construct.

### Protective efficacy of OV-HA and OV-HA-NP viruses intranasal IAV-S challenge

The protective efficacy of OV-HA and OV-HA-NP were evaluated upon intranasal challenge with IAV-S (after day 35 pi). Virus shedding was assessed in nasal secretions and viral load and pathology were evaluated in the lung. Nasal swabs were collected on days 0, 1, 3, and 7 post-challenge (pc) and IAV-S RNA levels were investigated in nasal secretions using real-time reverse transcriptase PCR (rRT-PCR). On day 1 pc, significantly lower IAV-S genome copy numbers – indicating reduced virus shedding – was detected in both OV-HA and OV-HA-NP immunized groups when compared to the control sham immunized group (Fig 6A). Only two animals (2/8) in the OV-HA-NP group were positive for viral RNA on day 1 pc. On 3 dpc, while all animals in control group (8/8) were positive and presented high genome copy numers of IAV-S in nasal secretions, only three animals (3/8) in OV-HA-NP were positive for viral RNA (Fig 6A). Notably, the amount of IAV-S RNA shed by OV-HA-NP-immunized animals were significantly lower than the amount shed by control or OV-HA immunized animals. It is also important to note that animals in OV-HA group had significantly lower level of viral RNA than control group on day 3 pi (Fig. 6A). On day 7 post-challenge, all animals (8/8) in the control sham-immunzied group were still shedding IAV-S in nasal secretions, while only two animals (2/8) in the OV-HA-immunized group were positive presenting low viral RNA copy numbers in nasal secretions. Notably, none of the animals in the OV-HA-NP-immunized group were shedding IAV-S in nasal sercetions on day 7 pi (Fig 6A). These results demonstrate that immunization with OV-HA and OV-HA-NP resulted in decreased virus shedding and shorter duration of virus shedding in nasal secretions following intranasal IAV-S challenge. Notably, these differences were more pronounced in OV-HA-NP-immunized animals.

**Figure 6.**
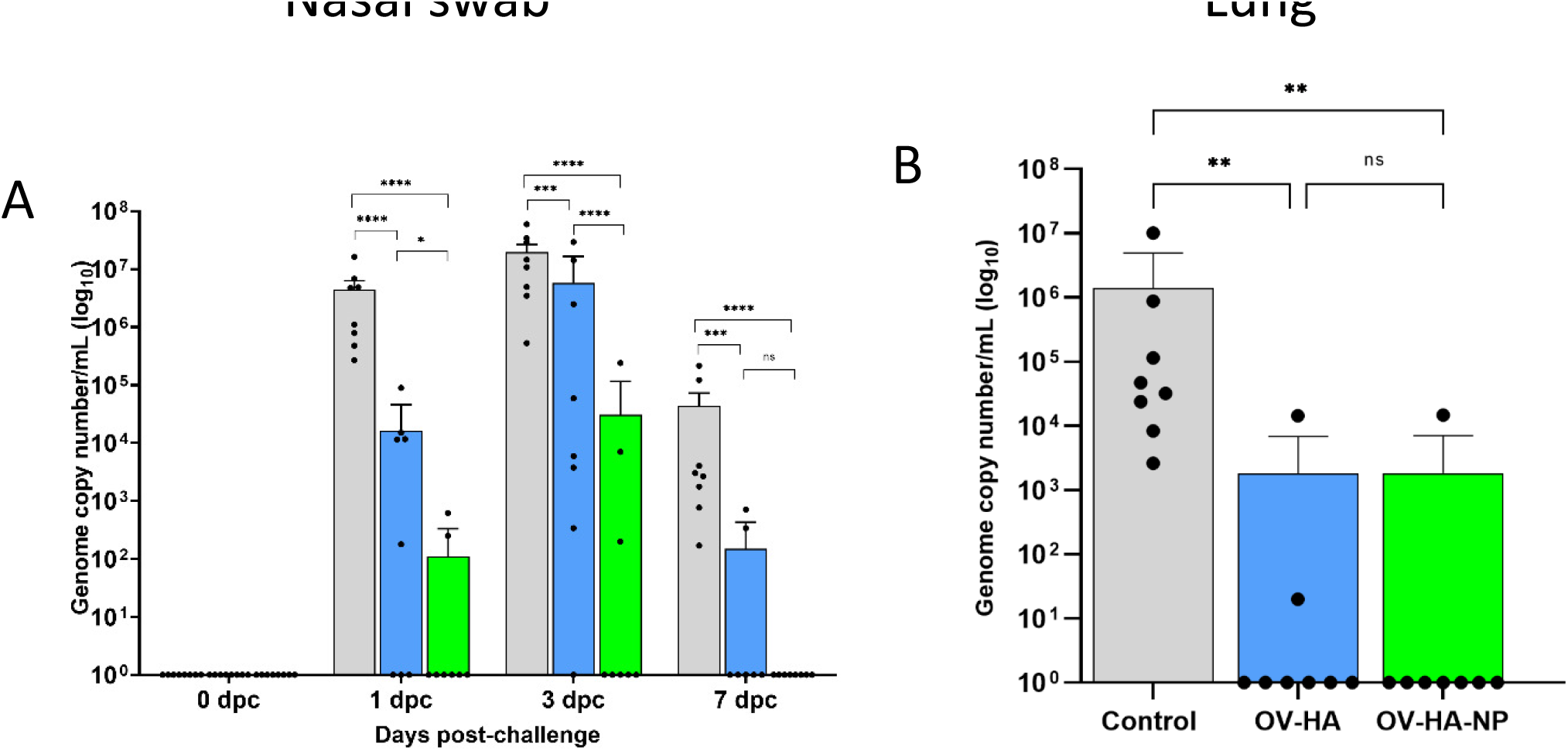
Protective efficacy of OV-HA and OV-HA-NP against IAV-S challenge. **(A)** IAV-S viral RNA shedding in the nasal swab was determined by RT-qPCR and expressed as log10 genome copy number per milliliter. **(B)** IAV-S viral load in the lung determined by RT-qPCR and expressed as log10 genome copy number per milliliter. Data represents group mean and error bars represent SEM. P-values: *\*P < 0*.*05, **P < 0*.*01, ***P < 0*.*001, ****P < 0*.*0001*.

Shedding of infectious IAV-S was also assessed in nasal secretions collected on days 0, 1, 3, and 7 post-challenge. Each sample was subjected to three blind passages in MDCK cells. An immunofluorescence assay using an IAV-S NP-specific monoclonal antibody was performed on the third passage to confirm isolation of IAV-S. On day 1 pc, 4 (50%) animals in the control group were positive for IAV-S, while none of the animals from the OV-HA and OV-HA-NP group were positive on VI (Table 2). On day 3 pc, 7 (87.5%) animals were positive in the sham-immunized control group; 3 (37.5%) animals were positive in OV-HA immunized group and 1 (12.5%) animal was positive in OV-HA-NP-immunized group. Statistical analysis confirmed that there was a significant difference in the number of IAV-S positive animals between control group and OV-HA-NP group on 3 dpc (P= 0.0101 Fisher’s exact test) (Table 2). IAV-S was not isolated from any of the animals on day 7 post-challenge (Table 2). These results indicate that both OV-HA and OV-HA-NP recombinants were able to reduce virus replication and shedding in the immunized animals. Importantly, detection of infectious virus in only one out of eight animals in OV-HA-NP groups highlingts the robust protection provided by immunization of pigs this recombinant virus.

**Table 2.**
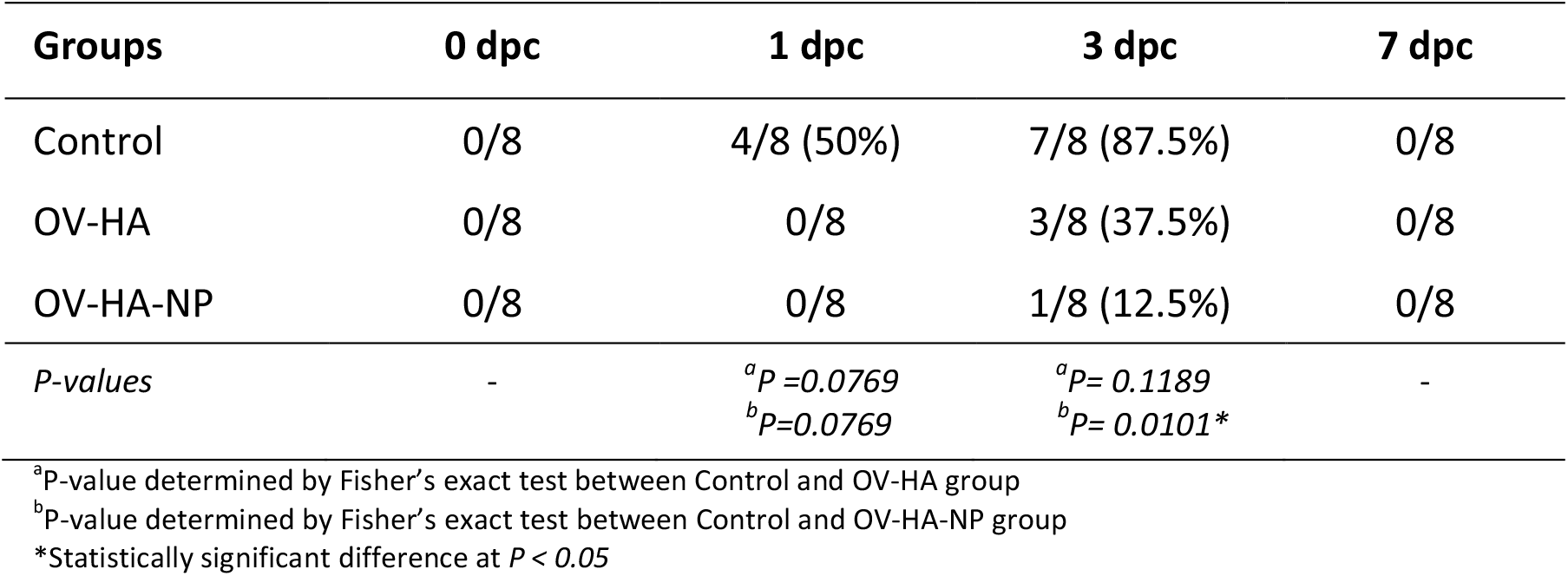
Virus isolation from the nasal swabs

| Groups | 0 dpc | 1 dpc | 3 dpc | 7 dpc |
| --- | --- | --- | --- | --- |
| Control | 0/8 | 4/8 (50%) | 7/8 (87.5%) | 0/8 |
| OV-HA | 0/8 | 0/8 | 3/8 (37.5%) | 0/8 |
| OV-HA-NP | 0/8 | 0/8 | 1/8 (12.5%) | 0/8 |
| *P-values* | - | <sup>a</sup>*P* = 0.0769<br><sup>b</sup>*P* = 0.0769 | <sup>a</sup>*P* = 0.1189<br><sup>b</sup>*P* = 0.0101 * | - |

<sup>a</sup>P-value determined by Fisher's exact test between Control and OV-HA group
<sup>b</sup>P-value determined by Fisher's exact test between Control and OV-HA-NP group
\*Statistically significant difference at *P* < 0.05

Viral load was assessed in the lung of control and immunized pigs on day 7 dpc by using rRT-PCR. While high amounts of IAV-S RNA were detected in the lung of animals in the control sham-immunized group, immunization with OV-HA or OV-HA-NP led to a marked decrease in viral load in the lung (Fig 6B). Notably, only one animal (1/8) in the OV-HA-NP group and two animals (2/8) in OV-HA group presented IAV-S RNA in lung, whereas all the animals in control group (8/8) were positive for IAV-S RNA. Significantly lower IAV-S RNA loads were detected in the lung of immunized animals when compared to control animals (Fig 6B)

In addition to viral loads pathological changes were also evaluated in the lung of all animals in the study. At necropsy, macroscopic lesions in the lung were characterized by a pathologist who was blinded to the experimental groups. A summary of the gross lung lesions is provided on Table 3. All animals in the control group presented characteristic plum-colored consolidated areas mostly on the cranioventral areas and interstitial pneumonia. Mild lobular consolidation and interstitial pneumonia was present in 2 animals in OV-HA group and 2 animals in OV-HA-NP group. As expected, the lesions were primarily observed in animals having relatively lower levels of neutralizing antibody titers (Table 3). No microscopic lesions were observed in any animals on day 7 post-challenge. Together these results indicate that immunization with ORFV-based vectors, especially with OV-HA-NP virus provided good protection against intranasal homologous IAV-S challenge in pigs.

**Table 3.** Pathological and serological findings post-IAV-S-challenge in immunized pigs.

| Control |  |  | OV-HA |  |  | OV-HA-NP |  |  |
| --- | --- | --- | --- | --- | --- | --- | --- | --- |
| Animal ID | Gross Lesions | VN Titer<sup>a</sup> | Animal ID | Gross Lesions | VN Titer<sup>a</sup> | Animal ID | Gross Lesions | VN Titer<sup>a</sup> |
| 22 | Lobular consolidation on the left cranioventral areas | <1:5 | 20 | Mild lobular consolidations on both right and left lung | 1:80 | 21 | No lesions | 1:5120 |
| 23 | Lobular consolidation present on ventral areas | <1:5 | 24 | No lesions | 1:320 | 25 | No lesions | 1:2560 |
| 29 | Lobular consolidation mostly present on cranioventral surface and interstitial inflammation on the left lobe | <1:5 | 26 | Mild lobular consolidation on both sides of the lung | 1:40 | 27 | No lesions | 1:640 |
| 30 | Lobular consolidation mostly present on cranioventral area | <1:5 | 33 | No lesions | 1:640 | 28 | No lesions | 1:1280 |
| 31 | Lobular consolidation mostly present on cranioventral area | <1:5 | 34 | No lesions | 1:2560 | 32 | No lesions | 1:1280 |
| 39 | Lobular consolidation mostly present on cranioventral area | <1:5 | 35 | No lesions | 1:1280 | 37 | Very mild lobular consolidation | 1:640 |
| 40 | Lobular consolidation mostly present on cranioventral area | <1:5 | 36 | No lesions | 1:640 | 38 | Congestion on apical lobe with mild interstitial pneumonia | 1:640 |
| 48 | Lobular consolidation on both sides of the cranioventral area | <1:5 | 49 | No lesions | 1:320 | 41 | No lesions | 1:2560 |

<sup>a</sup>VN titer measured on day 35 post-immunization.
All animals were terminated and examined on day 7 post-challenge and evaluated by a pathologist blinded to the study.

## Discussion

In this study we explored the potential of ORFV recombinants expressing the HA or both HA and NP proteins of IAV-S in providing protection against intranasal challenge infection in swine. Previous work from our group have shown that rational vector design by deleting well-characterized immunomodulatory genes of ORFV is useful in developing highly effective vaccine delivery platforms resulting in safe and highly immunogenic vaccine candidates. One of the well characterized ORFV IMPs is ORFV121, which encodes an NF-κB inhibitor that determines ORFV virulence and pathogenesis in the natural host (39). We have developed highly immunogenic vaccine candidates for porcine epidemic diarrhea virus (PEDV) and rabies virus (RabV) by inserting appropriate protective antigens (spike glycoprotein for PEDV; rabies glycoprotein for RabV) in the *ORFV121* gene locus. Given the immunogenicity and safety profile of the OV-PEDV-S and OV-RABV-G recombinant virus in swine, here we constructed an OV-HA recombinant by inserting the HA gene of IAV-S virus in ORFV121 locus.

Moreover, to potentially enhance T-cell immune response elicited by the vaccine we generated a second recombinant virus expressing both IAV-S NP and HA proteins. For this, another well-characterized ORFV IMP, the ORFV127 was selected as an insertion site for the NP gene. ORFV127 encodes a viral IL-10 homolog (15, 45), which is known to have anti-inflammatory and immunosuppressive activities that may favor immune evasion of the orf virus (46, 47). Most importantly, the protein encoded by ORFV127 is known to contribute to ORFV virulence in the natural host (48). Using this approach we tested the hypothesis that simultaneous deletion of two ORFV IMP genes ORFV121 and ORFV127 and concurrent insertion of two highly immunogenic protective antigens of IAV-S (HA and NP) would enhance the immunogenicity of the recombinant virus in swine and provide higher protective efficacy from IAV-S challenge. While the data presented here show that both recombinants OV-HA and OV-HA-NP induced robust immune response against IAV-S in pigs, the immunogenicity and protective efficacy of OV-HA-NP was indeed higher than that elicited by the OV-HA recombinant which substantiates our hypothesis. The increased protective efficacy seem to be related to stronger T-cell response elicited by immunization with the OV-HA-NP virus.

Following challenge infection, we observed interesting differences in the antibody response elicited by OV-HA and OV-HA-NP immunization. As expected, intranasal challenge with IAV-S in sham immunized pigs resulted in anamnestic VN response. Notably, in the immunized groups the VN antibody titers increased by a greater magnitude in the OV-HA group than in the OV-HA-NP group. In the OV-HA group, the geometric mean VN titers were 380.5 and 3319.9 on days 0 and 7 pc respectively, indicating a 9-fold increase in the VN titer after challenge. Whereas in OV-HA-NP group, the geometric mean VN titers were 1395.8 and 1810.19 on days 0 and 7 pc respectively, which is only a modest 1.2-fold increase in VN titers pc. Importantly, the VN titers in OV-HA-NP immunized group increased only in the two animals that had the lowest VN titer on day 0 pc. The VN titers in the remaining 6 animals in the OV-HA-NP group remained constant following challenge infection. These results demonstrate that immunization with the OV-HA-NP recombinant virus provided robust immune protection against intranasal IAV-S challenge, with most animals not seroconverting to the challenge virus.

The importance of T-cells in influenza virus clearance and their cross-reactive potential has been well documented (49, 50). In this context, CD4+ T-cells help with activation, differentiation and antibody production by virus-specific B cells (51). Additionally, CD4+ helper cells also play an important role in CD8+ cytotoxic T cell activation. Activated CD8+ cytotoxic T-cells function in virus clearance by killing infected cells (52). The NP protein of influenza virus is known to contain several immunologically dominant T-cell epitopes and it is the main antigen recognized by cytotoxic T-lymphocytes (CTL) during influenza A virus infections (42, 53–56). The Immune Epitope Database and Analysis Resource, a manually curated database of experimentally characterized immune epitopes, has recorded 248 T-cells epitopes for nucleoprotein (NP) of influenza virus. Given that NP is relatively conserved among influenza viruses, including in IAV-S, this protein has been one of the target viral antigens for the development of universal influenza vaccine candidates. Because of these important immunological properties, we have developed and evaluated the OV-HA-NP construct expressing both HA and NP proteins. We found that cell mediated immune responses were enhanced by co-delivery and expression of IAV-S HA and NP by OV-HA-NP in pigs when compared to OV-HA group. A significantly higher frequency of CD3+ T-cells proliferated and expressed IFN-γ upon re-stimulation with IAV-S in the OV-HA-NP-immunized group. Importanly, immunization with OV-HA-NP resulted in a increased frequency of CD3+/CD8+ T cells upon restimulation with IAV-S. While overall T-cell responses were higher in OV-HA-NP group, an increase in T-cell response was also seen in OV-HA group when compared to the sham immunized group, as evidenced by increase in IFN-γ secreting CD3+ T-cell population following antigen stimulation. This can be explained by the presence of several T-cells epitopes in the IAV HA protein, the majority of which have been identified as CD4+ T-cell epitopes (57, 58).

Depending upon the type of antigenic stimulation, CD4+ helper T-cell precursors (Th_o_) can either differentiate into Th1- or Th2-helper cells. Th1 cells secrete several cytokines including IFN-γ and IL-12 which help in cell mediated immunity, whereas Th2 cells secrete cytokines like IL-4, IL-6 which contribute to antibody mediated immunity (40, 59). Importantly, IgG isotype expression is also controlled by the different cytokines (60, 61). In pigs, IFN-γ enhances production of IgG2 isotype and hence this IgG isotype is considered to be associated with Th1 immune response. On the other hand, cytokines like IL-4, IL-10 induce secretion of IgG1 and are known to be associated with Th2 immune response (62). Thus, the ratio of IgG1:IgG2 can be used to infer Th1/Th2 bias in response to vaccination. In this study, we found a higher level of IgG1 in pigs immunized with OV-HA recombinant (IgG1:IgG2 >1, Th2 bias), which suggests that the protection may have been mostly antibody-mediated in this group. Conversely, in OV-HA-NP group, the levels of IgG2 were higher (IgG1:IgG2 <1, Th1 bias), which suggests a bias towards cell-mediated immunity in this group. Given that NP protein is known to induce cell-mediated immunity, it would be safe to assume that this Th1 bias might be due to NP protein present in the OV-HA-NP recombinant.

This study further demonstrates the use of ORFV as a vaccine delivery platform in swine. The study also shows that two ORFV IMP encoding genes (ORFV121 and ORFV127) can be deleted simulateously from the virus genome to efficiently delivery at least two viral antigens in swine. One of the advantages of ORFV-based vectors is that same vector can be used repeatedly for prime-boost regimens. This is important because pre-existing immunity precludes the use of many vector platforms for vaccine delivery. The humoral immune response data presented here shows that a boost effect was induced after second immunization. In fact, previous findings from our lab show that similar effect can be observed even after three immunizations with ORFV. The recombinant HA and NP protein used in this study share 95% animo acid identity with the HA and NP protein of the challenge virus. In future, we plan to use the HA gene from other IAV-S subtypes to develop multivalent vaccine candidates and evaluate heterosubtypic protection. The analysis of secretory IgA immune response, which play an important role in providing mucosal immune response is lacking in this study. Future studies involving detailed analysis of mucosal immune response elicited by ORFV-based constructs and challenge infection with heterologous IAV strains are warranted. Nonetheless, results presented here demonstrate that ORFV-based vectors can be important tools to develop improved vaccine candidates to effectively control IAV-S infections in swine.

## Material and methods

### Cells and viruses

Primary ovine turbinate cells (OFTu), Madin-Darby canine kidney cells (MDCK) and swine turbinate cells (STU) were cultured at 37 °C with 5% CO_2_ in minimum essential medium (MEM) supplemented with 10% FBS, 2 mM L-glutamine and containing streptomycin (100 µg/mL), penicillin (100 U/mL and gentamycin (50 µg/mL).

The ORFV strain IA82 (OV-IA82; kindly provided by Dr. Daniel Rock at University of Illinoir Urbana-Champaign), was used as the parental virus to construct the recombinants and in all the experiments involving the use of wild-type ORFV. Wild-type and recombinant ORFV viruses were amplified in OFTu cells. Swine influenza virus H1N1 A/Swine/OH/24366/2007 (H1N1), kindly provided by Gourapura Lab was used for virus challenge, virus neutralization assay, hemagglutination inhibition (HI), and as a coating antigen for whole virus ELISA. The H1N1 A/Swine/OH/24366/2007 (H1N1) virus was propagated in MDCK cells using DMEM containing TPCK-treated trypsin (2 µg/mL) and 25 mM HEPES buffer.

### Generations of recombination plasmids

To insert the heterologous IAV-S gene in the ORFV121 locus, a recombination plasmid containing right and left flanking sequences of the ORFV121 gene were inserted into pUC57 plasmid. The HA gene of swine influneza virus, A/SW/OH/511445/2007 (OH7) (GenBank : EU604689) (63) was inserted between the ORFV121 flanking sequence in the pUC57 plasmid. The HA gene was condon optimized for swine species (GenScript). The HA gene was cloned under the vaccinia virus (VACV) I1L promoter (5’-TATTTAAAAGTTGTTTGGTGAACTTAAATGG – 3’) (43) and a flag-tag epitope (DYKDDDK) was fused to the amino terminus of the HA gene to detect its expression. The gene encoding green fluorescent protein (GFP) was inserted downstream of HA gene and used as a selection marker for recombinant virus purification. The GFP sequence was flanked by *loxp* sequences 5’-ATAACTTCGTATAATGTATACTATACGAAGTTAT-3’ to allow for removal of GFP by Cre recombinase following recombinant virus purification. This recombination cassette was named pUC57-121LR-SIV-HA-loxp-GFP (Fig 1A).

Similarly, another recombination cassette was generated to insert NP gene of IAV-S into the ORFV ORFV127 locus. A recombination cassette for ORFV127 was constructed as describe above with the ORFV127 left and right flanking regions being cloned into the pUC57-LoxP-GFP plasmid (pUC57-127LR-LoxP-GFP. The nucleoprotein (NP) gene of swine influenza virus, A/SW/OH/511445/2007 (OH7) (GenBank: EU604694) (63) was inserted between ORFV127 left and right flanks. The NP gene was cloned under the VACV vv7.5 promoter (44) and the HA epitope tag sequence (YPYDVPDYA) was fused at the amino terminus of the NP protein to detect its expression by the recombinant virus. In addition, an eukaryotic Kozak consensus sequence 5’-gccaaccATGg-3’ (64), where ATG refers to the start codon of the NP gene, was added immediately downstream of vv7.5 promoter. This recombination cassette was named pUC57-127LR-SIV-NP-loxp-GFP (Fig 1B).

### Generation of recombinant OV-HA and OV-HA-NP viruses

The HA gene of IAV-S was inserted into the ORFV121 locus of the ORFV genome by homologous recombination. Briefly, OFTu cells cultured in 6-well plate were infected with OV-IA82 with a multiplicity of infection (MOI) of 1. Three hours later, the infected cells were transfected with 2 µg of pUC57-121LR-SIV-HA-loxp-GFP using Lipofectamine 3000 according to the manufacturer’s instruction (Invitrogen, catalog no: L3000-075). At 48 hours post-infection/transfection cell cultured were harvested, subjected to three freeze-and-thaw cycles. The ORFV recombinant expressing IAV-S HA was purified using plaque assay by selecting viral foci expressing GFP. After several rounds of plaque purification, the presence of HA gene and absence of ORFV121 gene was confirmed by PCR as described before (22, 24) and the insertion and integrity of the whole genome sequence of the recombinant was confirmed sequencing using Nextera XT DNA library preparation following by sequencing on the Illumina Mi-Seq sequencing platform. Once the purified recombinant virus was obtained, the GFP selection gene was removed by using Cre recombinase treatment as described below. This recombinant is referred to as OV-HA throughout this manuscript.

Similarly, double gene expression vector containing the IAV-S HA and NP genes in ORFV121 and the ORFV127 gene loci (48), respectively was generated by homologous recombination. Both ORFV121 and ORFV127 are virulence determinants that contribute to ORFV IA-82 virulence in the natural host (39, 48). For this, infection/transfection was performed by infecting OFTu cells with the OV-HA recombinant virus and transfecting with pUC57-127LR-SIV-NP-loxp-GFP plasmid. The recombinant virus was purified using plaque assay as described above and following purification the GFP reporter gene was removed using the Cre recombinase treatment described below. The resulting recombinant ORFV vector expressing the HA and NP gene is referred to as OV-HA-NP in this manuscript.

The Cre/loxP recombination system was used to remove the GFP reporter gene from the OV-HA or OV-HA-NP recombinants. A plasmid pBS185 CMV-Cre, carrying the cre gene under the hCMV promoter was a kind gift from Brian Sauer (65) (Addgene catalog number : 11916). OFTu cells were plated in a 24-well plate and 24h later transfected with 500 ng of the pBS185-CMV-Cre plasmid using Lipofectamine 3000 (Invitrogen, catalog num: L3000-075) according to the manufacturer’s instructions. Approximately 24h after transfrections, cells were infected with ∼ 1 MOI of the plaque purified recombinant viruses (OV-HA-GFP or OV-HA-NP-GFP). Approximately 48 h post-infection, the cre recombinase treated recombinant viruses were harvested and subjected to a second round of Cre treatment as described above. Following cre recombinase treatment, two to three rounds of plaque assays were performed to select foci lacking GFP expression and to obtain reporterless OV-HA or OV-HA-NP recombinant viruses. Following markerless virus selection complete genome sequencing was performed to determine the integrity of ORFV and IAV-S sequences in the recombinant OV-HA and OV-HA-NP viruses.

### Growth curves

Replication kinetics of OV-HA and OV-HA-NP recombinant viruses were assessed *in vitro* in OFTu and STU cells. Briefly, OFTu and STU cells cultured in 12-well plates were inoculated with OV-HA or OV-HA-NP with a multiplicity of infection (MOI) of 0.1 (multistep growth curve) or 10 (single-step growth curve) and harvested at 6, 12, 24, 48, 72 hours post-infection (hpi). Virus titers in cell lysates and supernatants were determined on each time point using Sperman and Karber’s method and expressed as tissue culture infectious dose 50 (TCID_50_) per milliliter (66).

### Immunofluorescence

Immunofluorescence assay (IFA) was used to assess expression of the heterologous proteins by the OV-HA or the OV-HA-NP viruses as described previously (67). Briefly, OFTu cells were inoculated with each recombinant virus (MOI of 1) and fixed with 3.7% formaldehyde at 48 hours pi. Then, cells were permeabilized with 0.2% PBS-Triton X-100 for 10 min at room temperature. Another set of samples which were not permealized were also tested side-by-side to compare the expression pattern between permeabilized and non-permeabilied cells. Flag-tag specific mouse antibody (Genscript, catalog no: A100187) and HA-tag specific rabbit antibody (Cell Signaling, catalog no: 3724S) were used as primary antibody to detect HA and NP protein respectively. Then, cells were incubated with Alexa fluor 594 goat anti-mouse IgG (H+L) secondary antibody (Invitrogen, catalog no: A11005) or Alexa fluor 488 goat anti-rabbit IgG antibody and cells were observed under fluorescence microscope.

### Animal immunization and challenge studies

The immunogenicity of the two recombinant viruses (OV-HA and OV-HA-NP) was evaluated in 3-week old high-health pigs. A summary of experimental design is presented in Table 1. Twenty-four pigs, seronegative for IAV-S, were randomly allocated into three experimental groups as follows: Group 1, sham immunized (n=8); Group 2, OV-HA immunized (n=8); Group 3, OV-HA-NP immunized (n=8). Immunization was performed by intramuscular injection of 2 ml of a virus suspension containing 10^7^ TCID_50_/mL in MEM. All animals were immunized on day 0 and received a booster immunization on day 21 post-immunization. All animals were challenged intranasally on day 35 post-immunization with 5 mL virus inoculum containing 6 × 10^6^ TCID_50_ of H1N1 A/Swine/OH/24366/2007 (H1N1) (68) per animal. Animals were monitored daily for clinical signs of IAV-S. Serum and PBMC samples were collected on days 0, 7, 14, 21, 28, 35, 38 and 42 days post-immunization. Nasal swabs were collected on days 0, 1, 3, 7 post-challenge. The experiment was terminated on day 42 post-immunization or 7 days post-challenge. Whole lung as a unit were collected from euthanized animals during necropsy and examined grossly for pathologic changes by a pathologist blided to study groups. Animal immunization challenge studies were conducted at South Dakota State University (SDSU) Animal Resource Wing (ARW), following the guidelines and protocols approved by the SDSU Institutional Animal Care and Use Committee (IACUC approval no. 17-018A)

### Virus neutralization (VN) assay

Virus neutralization titer in the serum samples were determined as described previously (69). Briefly, serum samples were heat inactivated for 30 minutes at 56 °C. Two-fold serial dilutions of serum were incubated with 200 TCID_50_ of IAV-S, A/Swine/OH/24366/2007 (H1N1), at 37 °C for 1 hour. This virus-serum complex was then transferred to a 96-well plate pre-seeded with MDCK cells 24 h earlier. After 1 hour of adsorption, virus-serum complex was removed and fresh DMEM containing 2 µg/mL of TPCK-treated trypsin was added to the cells. After 48-hour incubation at 37 °C, cells were fixed with 80% acetone. Virus positive MDCK cells were detected by immunofluorescence assay using a mouse monoclonal antibody targeting nucleoprotein (NP) of influenza virus (IAV-NP HB-65 mAb; kindly provided by Drs. Eric Nelson and Steve Lawson at SDSU). The virus neutralization titer was defined as the reciprocal of the highest dilution of serum where there was complete inhibition of infection/replication as evidenced by absence of fluorescent foci. Appropriate positive and negative control samples were included in all the plates.

### Hemagglutination inhibition (HI) assay

HI assay was performed according to the method descrined previously (69).Briefly, serial 2-fold dilution (starting dilution 1:4) were prepared in PBS. Then 4 HA units of H1N1 A/Swine/OH/24366/2007 virus was added to the serum dilutions and incubated at room temperature for 1 hour. A solution (in PBS) of turkey red blood cells (containing 0.5% RBC) were added to the wells and allowed to settle. The HI titer was calculated as the reciprocal of the highest dilution of sera that inhibited hemagglutination of turkey RBC.

### Real-time reverse transcriptase PCR (rRT-PCR)

Virus shedding in nasasl secretions and viral load in lungs was evaluated by rRT-PCR. Lung tissues were homogenized using tissue homogenizer by adding 10 mL of DMEM in 1 g of lung tissue. Viral nucleic acid was extracted from the nasal swabs and lung tissue homogenates using the MagMax Viral RNA/DNA isolation Kit (Life Technologies). The rRT-PCR tests were performed at Animal Disease Research and Diagnostic Lab (ADRDL), SDSU, SD. Genome copy numbers per milliliter were determined based on the relative standard curve derived from four-parameter logistic regression analysis (*R-square=0*.*9928, Root mean square error (RMSE)=1*.*0012*).

### Virus isolation

Virus isolation was performed on the nasal swabs collected on day 0, 1, 3, and 7 post-challenge. Nasal swabs were filtered through a 0.22-micron filter and mixed with DMEM containing 2 µg/mL of TPCK-treated trypsin in 1:1 ratio. Then, 250 µL of this inoculum was added to 24-well plate containing MDCK cells. The cells were incubated for 1 hour at 37 °C. After 1 hour adsorption, 250 µL of DMEM was added to the wells and plate was incubated for 48 hours. After 48 hours, cell lysate was harvested, and two more blind serial passages were performed. After the third passage, the supernatant was collected, and the cells were fixed with 80% acetone. Immunofluorescence assay (IFA) was performed using IAV-NP mAb (IAV-NP HB-65) as primary antibody and Alexa fluor 594 goat anti-mouse antibody as secondary antibody (Invitrogen, catalog no: A11005). SIV infected cells were identified based on the presence of fluorescent foci.

### ELISA

IAV-S-specific IgG, IgG1a and IgG2a immune response elicited by immunization with OV-HA or OV-HA-NP were assessed by whole virus ELISA. The antigen for coating the ELISA plates was prepared as described previosly (70) with some modifications. Briefly, ultra-centrifugation of virus culuture supernatant and the virus pellet in 30% sucrose cushion gradient were performed using Optima-L 100K ultracentrifuge (Beckman Coulter) at 18,000 RPM for 1.5 hours. The virus pellet was resuspended in DMEM and UV inactivation of the virus was carried out using CL1000 UV crosslinker. Determination of the optimal coating antigen concentration and dilution of secondary antibodies were carried out by checkerboard titration.

To detect IAV-S specific total IgG, Immulon 1B ELISA plates (ThermoFisher Scientific, catalog no: 3355) were coated with 250 ng/well of concentrated and UV inactivated IAV-S virus and incubated at 37 °C for 2 hours. Then plates were washed three times with PBST (1X PBS with 0.5% Tween-20) and blocked with 200 µL/well of blocking solution (5% milk in PBST) and incubated overnight at 4 °C. Then, the plates were washed three times with PBST. Serum samples diluted in blocking solution at the dilution of 1:100 was added, and the plates were incubated for 1 hr at room temperature (RT). After, three washes with PBST, 100 µL of biotinylated anti-pig IgG antibody (Bethyl, catalog no: A100-104) diluted in blocking buffer (1:4000) was added to the plate and incubated for 1 hr at RT. Following three washes, HRP-conjugated streptavidin (Thermo Scientific, catalog no: 21136) diluted in blocking solution (1:4000) was added to plates and incubated for 1 hr at RT. Plates were washed again for three times with PBST and 100 µL/well of 3,3′,5,5′-tetramethylbenzidine (TMB) substrate was added to the plates (KPL, catalog no: 5120-0047). Finally, the colorimetric reaction was stopped by adding 100 µL 1N HCl solution per well. Optical density (OD) values were measured at 450 nm using a microplate reader. Cut-off value was determined as mean OD of negative serum samples plus three times of standard deviation (mean + 3SD).

Isotype ELISA were performed on the serum samples collected on day 35 post-immunization. For isotype ELISA, mouse anti-pig IgG1 (Biorad, catalog no: MCA635GA) and mouse anti-pig IgG2 antibody (Biorad, catalog no: MCA636GA) were used as secondary antibodies and plates were incubated with biotinylated anti-mouse antibody (KPL, catalog no: 5260-0048) before incubating with streptavidin-HRP antibody. Endpoint titer ELISA using serial two-fold serial dilutions of serum samples were performed to determine endpoint titer of SIV-specific IgG1 and IgG2 antibody levels in the serum samples. Other procedures were similar to the total IgG ELISA as described above.

### Flow-cytometry

IAV-S-specific T-cell response elicited by ORFV recombinants was evaluated by an intracellular cytokine staining (ICS) assay for interferon gamma (IFN-γ) and T-cell proliferation assay. For IFN-γ expression assay, cryopreserved PBMCs collected on day 35 post-immunization (0 dpc) were thawed and seeded at a density of 5 × 10^5^ cells/well in 96-well plate. Cells were stimulated with UV inactivated IAV-S at MOI of 1. Additionally, cells were stimulated with concanavalin (ConA: 2 μg/ml) (Sigma, catalog no: C0412) plus phytohemagglutinin (PHA: 5 μg/ml) (Sigma, catalog no: 61764) as positive control and cRPMI (RPMI with 10% FBS) was added to the negative control wells. Protein transport inhibitor, Brefeldin A (BD Biosciences, catalog no: 555029), was added 6 hours after stimulation and the cells were incubated for 12 hours prior to flow cytometric analysis. For the proliferation assay, PBMCs (35 dpi) were stained with 2.5 μM carboxyfluorescin succinimidyl ester (CFSE; in PBS) (BD Horizon, catalog no: 565082). CFSE stained cells were seeded at a density of 5 × 10^5^ cells/well in 96-well plate. The cells were stimulated as described above. After stimulation, the cells were incubated for 4 days at 37°C with 5% CO_2_ prior to staining. Antibodies used for immunostaning the cells were : CD3+ (Mouse anti-pig CD3ε Alexa Fluor 647; BD Pharmingen, catalog no: 561476), CD4+ (Primary antibody: Mouse anti-pig CD4, Monoclonal Antibody Center (WSU), catalog no: 74-12-4; secondary antibody: Goat anti-mouse IgG2b PE/Cy7, Southern Biotech, catalog no: 1090-17), CD8+ (Primary antibody: Mouse anti-pig CD8α, Monoclonal Antibody Center (WSU), catalog no: 76-2-11; secondary antibody: Goat anti-mouse IgG2a FitC, Southern Biotech, catalog no: 1080-02), IFN-γ (Anti-pig IFN-γ PE, BD Pharmingen, catalog no: 559812. The stained cells were analyzed using Attune NxT flow-cytometer. Results were corrected for background proliferation by subtracting mock-stimulated proliferation from the frequency of cells that responded under inactivated SIV stimulation. The percentage of responding cells was calculated as the percentage of total T cells (live CD3+ cells).

### Statistical analysis

Statistical analysis was performed using Graphpad Prism software. The normality of the data was tested using Shapiro-Wilk test. Comparison of means between the groups was done using two-way ANOVA for normal data or Kruskal Wallis test for non-normal data. Pairwise comparison was done using Tukey multiple comparison test. P value of less than 0.05 was considered significant. Flow cytometry data was analyzed using Flow Jo software.

## Acknowldgement

We thank the Animal Resource Wing (ARW) SDSU for their assistance in animal experiments. We thank the Cornell BRC Flow Cytometry Facility at the Cornell Institute of Biotechnology for the use of flow cytometers for data acquisition. We also would like to thank Bishwas Sharma, Maureen Hoch Vieira Fernandes, Jessica Caroline Gomes Noll, Gabriela Mansano do Nascimento, and Steve Lawson for their help with sample collection. This work was supported by AFRI Foundational and Applied Science Program (grant no. 2017-67015-32034/project accession no. NYCV478904) from the USDA National Institute of Food and Agriculture.

